# Reconciling ecology and evolutionary game theory or ‘When not to think cooperation’

**DOI:** 10.1101/2024.07.10.602961

**Authors:** Corina E. Tarnita, Arne Traulsen

## Abstract

Evolutionary game theory (EGT)—overwhelmingly employed today for the study of cooperation in a variety of systems, from microbes to cancer and from insect to human societies—started with the seminal 1973 paper by John Maynard Smith and George Price [1], in which they probed the logic of limited war in animal conflict. If fighting was essential to get access to mates and territory, then why did fights rarely lead to serious injury? Maynard Smith and Price developed game theory to show that limited war can be selected at the individual level. Owing to the explanatory potential of this first paper, and enabled by the elegant and powerful machinery of the soon-to-be-developed replicator dynamics [2, 3], EGT took off at an accelerated pace and began to shape expectations across systems and scales. But, even as it expanded its reach from animals to microbes [4–8] and from microbes to cancer [9–11], the field did not revisit a fundamental assumption of that first paper, which subsequently got weaved into the very fabric of the framework—that individual differences in reproduction are determined only by payoff from the game (i.e. in isolation, all individuals, regardless of strategy, were assumed to have identical intrinsic growth rates). Here, we argue that this original assumption substantially limits the scope of EGT. But, because it is not explicitly presented as a caveat, predictions of EGT have been empirically tested broadly across real systems, where the intrinsic growth rates are generally not equal. That has, unsurprisingly, led to puzzling findings and contentious debates [7, 12–15]. Flagging the high potential for confusion to arise from applications of EGT to empirical systems that it is not designed to study and suggesting a way forward constitute our main motivation for this work. In the process, we reestablish a dialog with ecology that can be fruitful both ways, e.g., by providing a so-far-elusive explanation for how diverse ecological communities can assemble evolutionarily.

---

Game theory as a mathematical discipline was founded by John von Neumann at the beginning of the 20th century and established as a relevant perspective on economic behavior via the book *Theory of Games and Economic Behavior* that von Neumann co-authored in 1944 with the economist Oskar Morgenstern [16]. It was not long before biologists realized that a game theoretic perspective might also be fruitful in biology. By the early 1960s, Lewontin suggested that “the modern theory of games may be useful in finding exact answers to problems of evolution not covered by the theory of population genetics”[17] and Slobodkin contemplated an analogy between his experimental findings on interspecific interactions and the result of species playing games with the environment [18]. These early attempts showed great foresight and were, in some ways, richer in their interpretation of potential evolutionary games than what ultimately became EGT, but they relied either on imprecise verbal analogies or on unwieldy mathematical formulations. It would be another decade before Maynard Smith and Price [1] achieved the trifecta of (i) a compelling and elusive question, with (ii) a simple, tractable and biologically meaningful game theoretic formulation that yielded (iii) an insightful and biologically coherent answer.

Prior to Maynard Smith and Price [1], explanations for limited conflict were grounded in a ‘species-selection’ view whereby total war would be bad for the species. Maynard Smith and Price explored several strategies that males of the same species might employ during conflict, but the two that would become most famous for illustrating the power of the new EGT framework were Hawk (aggressive males that fight to death or until serious injury) and Dove (pacifist males that backed down from conflict; originally called ‘Mouse’[1]). The game matrix

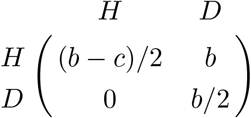

captures with fewest parameters the expected payoff obtained in a pairwise encounter: two Doves would split the benefit *b >* 0, a Dove encountering a Hawk would retreat, getting payoff 0 and yielding the entire benefit to the Hawk, and two Hawks would fight a total war resulting in an expected payoff of (*b* − *c*)*/*2 *<* 0, where *c > b >* 0 to reflect the high cost of total war. Under the assumption that the only difference in fitness between males of the same species is the fighting strategy employed, Maynard Smith and Price investigated whether either of these strategies would be evolutionarily stable (ESS). For a strategy to be evolutionarily stable, it must be resistant to invasion by rare mutants [1]. Applying this concept to the game matrix above, the Hawk strategy is ESS if its payoff playing against itself exceeds the payoff that a mutant Dove strategy gets from playing against the resident Hawk. This analysis revealed that neither Hawk nor Dove is ESS and that the higher the cost of total war, the worse a population of all Hawk fares.

From these results one can intuit that, since neither strategy is ESS, invaders of either type will grow into a resident population of the other type, but not indefinitely. The population should thus equilibrate at a mix of the two strategies, with the proportion of Hawk decreasing with the magnitude of the cost relative to the benefit. That sufficed for Maynard Smith and Price to demonstrate that selection at the individual level does not favor a full Hawk population (i.e. total war), but a mix of some aggressive individuals with mostly pacifist ones. Because this paper revealed such enormous potential for the “modern theory of games”[17] to bring clarity and alternative hypotheses to elusive questions in animal behavior, mathematical biologists [2, 3, 19] quickly began to develop a dynamical theory that allowed the exploration of the full strategy space while recapitulating the outcomes of the ESS approach in the special case of the homogeneous equilibria. Operating still under the original assumption that individual differences in reproduction are determined only by payoff from the game, the dynamical equation giving the change in the frequency of strategy *i* with time took the simplest form of an exponential growth or decay, 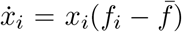, where—much as R.A. Fisher had proposed earlier for constant selection [20]—the growth rate was given by the difference between *f*_*i*_, the fitness that an individual using strategy *i* derives from the game (which depends on the matrix entries and on the frequency of each strategy in the population), and, 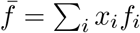 the average population fitness.

These equations became known as the **replicator dynamics** but, though dynamically richer than the original ESS analysis, they too were built on the assumption that the only difference in the reproductive ability of different strategies comes from differential payoffs within the game itself [21]. But while that may have been a reasonable assumption in snakes, mule deer, and Arabian oryx, where Maynard Smith and Price first situated their thinking, it is far from clear that this assumption holds true generally for microbes or cells. Nonetheless, this assumption has been embedded so deeply into the fabric of EGT that it has stopped being explicitly acknowledged as a simplification or caveat, which has lead to applications of EGT and its predictions to systems where this original assumption might not hold [12, 13]. All this raises two fundamental questions: if we did not make this assumption, what would be the simplest extension of replicator dynamics that also allows for intrinsic differences between types/strategies? And, importantly, would the predictions of such an extended framework differ from those of EGT when there are intrinsic differences between the growth rates of the two types?

Such a framework, turns out, already exists and is in fact much older than evolutionary game theory: it is given by the **generalized Lotka-Volterra equations**, where the growth rate of each type is determined not just by interactions with itself and others, but also by an intrinsic growth rate *r* (i.e. a type-specific growth rate when rare, in the absence of any interactions). The generalized Lotka-Volterra equations thus capture the dynamics of absolute abundances, *n*_*i*_, as 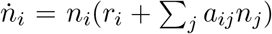, where *a*_*ij*_ is the density-dependent effect^*^ of type *j* on type *i*. Although the Lotka-Volterra equations were written by ecologists to think about species interactions, there is nothing in the equations themselves that precludes their interpretation in terms of types in general.

## Does the original Maynard Smith and Price assumption limit EGT substantially?

That the generalized Lotka-Volterra (LV) is precisely the simplest extension of replicator dynamics (RD) that accounts for intrinsic growth rate differences between types can be easily seen from the equations. For simplicity of exposition we will restrict ourselves to two types, but the relationship between the two frameworks derived below in Eq. (3) can be generalized to, and raises qualitatively the same issues for any number of types (see Appendix A).

For two types interacting according to the general matrix

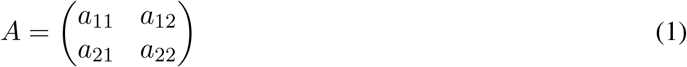

where *a*_*ij*_ is the effect that a type *j* individual has on the growth rate of a type *i* individual, we can write the **Lotka-Volterra equations** to track the dynamics of their abundances:

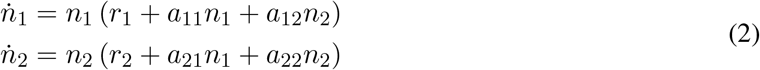

From the dynamics of abundances, we can use the quotient rule to derive the dynamics of relative frequencies, *x*_1_ = *n*_1_*/*(*n*_1_ + *n*_2_) and *x*_2_ = *n*_2_*/*(*n*_1_ + *n*_2_), and obtain:

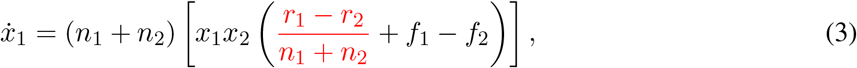

where we have denoted *f*_1_ := *a*_11_*x*_1_ + *a*_12_*x*_2_ and *f*_2_ := *a*_21_*x*_1_ + *a*_22_*x*_2_ (see Appendix A for the step-by-step derivation). The equation for *x*_2_ can be similarly derived, but it is redundant since *x*_2_ = 1 − *x*_1_.

If the two types have the same intrinsic growth rates (i.e. *r*_1_ = *r*_2_) or the population grows without bounds (i.e. *n*_1_ + *n*_2_ → ∞), then the red term in Eq. (3) becomes zero and the equation is dynamically identical (up to a rescaling of time by the positive factor *n*_1_ + *n*_2_) to the equation

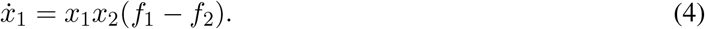

This equation is precisely the **replicator equation** 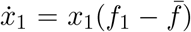 for two types playing the game given by matrix *A* in (1), where *f*_1_ = *a*_11_*x*_1_ + *a*_12_*x*_2_ is interpreted as the fitness of an individual of type 1 and 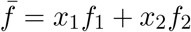 is the average population fitness^†^.

This derivation—and its general form for *m* types shown in Appendix A—recapitulates an equivalence established many decades ago [2, 22–24] between the *m*-type Lotka-Volterra system with identical growth rates and the replicator dynamics for an *m*-type game. But it also explicitly highlights the conditions that are both necessary and sufficient for the equivalence, thereby allowing us to engage in depth with the scenarios where the two conditions are violated. Specifically, when the system is bounded and the two types have different intrinsic growth rates, the red term in Eq. (3) can be non-negligible, and there is no general equivalence between RD and LV, because the dynamics of frequencies cannot be separated from the dynamics of abundances in Eq. (3). A natural reaction is that, even when the system is bounded, if it is very large (e.g., as is the case in microbial populations) the problematic term in Eq.(3) might be negligible. This intuition, however, is incorrect because what matters is the magnitude of that term at the point where *f*_1_ = *f*_2_ and, there, the red term can still be non-negligible when *r*_1_ ≠ *r*_2_. This can substantially change the coexistence fixed point and even its stability.

In other words, if the two types have different intrinsic growth rates, the outcome of their dynamics might not be predicted by the replicator equation^‡^, which means that the original assumption of Maynard Smith and Price—that fitness differences between individuals arise only from the game—does substantially limit the scope of EGT. Applying its predictions broadly to empirical systems that it was not designed to study has high potential for confusion^§^.

### What can go wrong?

To catalog the possible discrepancies, we contrast the predictions of the two frameworks in a bounded system with and without equal intrinsic growth rates (Fig. 1). For the system to be bounded, either type in isolation must be bounded (i.e. *a*_*ii*_ *<* 0 for *i* = 1, 2) and the system with both types present must also be bounded (i.e. *a*_12_*a*_21_ *< a*_11_*a*_22_). Without loss of generality, we assume that type 2 experiences the stronger intra-type competition, i.e. *a*_11_ *> a*_22_ (equivalently, that type 1 gets a higher payoff against itself than type 2 does against itself). The black and white plot in Fig. 1 illustrates the three main ecological interactions that matrix (1) captures depending on the signs of the interspecific parameters (*a*_12_, *a*_21_): mutualism (+, +), predator-prey or parasitism (+, − or *vice versa*), and competition (−, −). This ecological classification—although textbook in guiding ecological thinking [25]—does not directly reveal the dynamical outcome of the interaction. For one, this classification itself ignores the intrinsic growth rates. But even for identical growth rates, the same type of ecological interaction can have different outcomes depending on the values of the interaction terms: e.g., predator-prey dynamics can result either in coexistence or in the extinction of prey. A century of analysis of the Lotka-Volterra equations has yielded a thorough understanding of their dynamical outcomes. We contrast these with the predictions of replicator dynamics.

**Figure 1:**
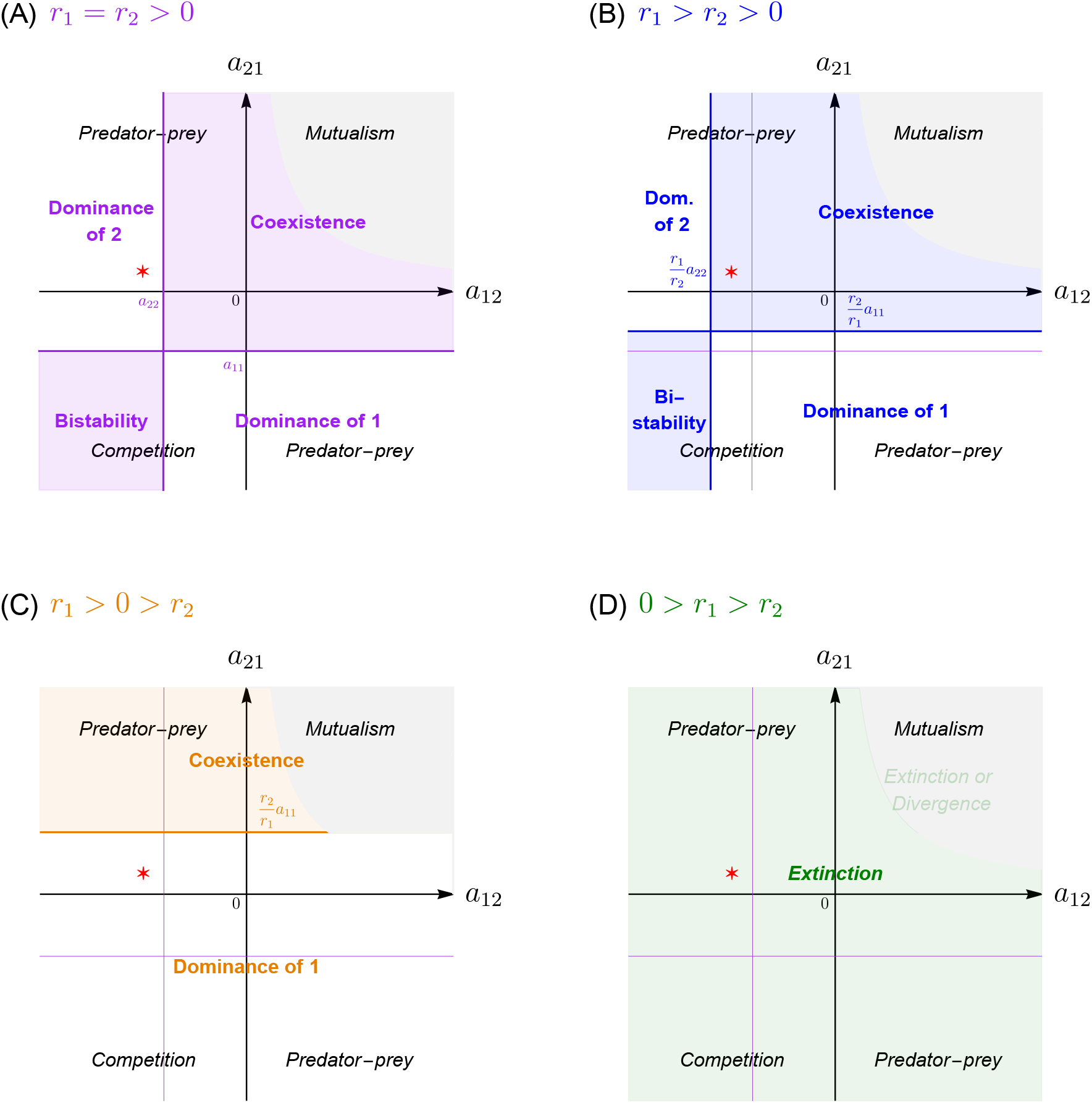
Ecology & Evolutionary Game Theory. Relationship between ecological classification of interaction type (in black and white, A-D), game theoretical classification of the dynamics under the assumption that *r*_1_ = *r*_2_ (in purple, A-D), and correct dynamical classification when *r*_1_ ≠ *r*_2_ (B, in blue; C, in orange; D in green). In the gray areas, *a*_12_*a*_21_ *> a*_11_*a*_22_ and system is unbounded. **(A)** When *r*_1_ ≠ *r*_2_ *>* 0, the replicator dynamics (RD) predicts accurately the four possible dynamical outcomes of the ecological interactions (which differ from the ecological classification into interaction types). **(B-D)** When *r*_1_ = *r*_2_, the RD does not accurately predict the outcome of the ecological interactions. Correctly predicting dynamics using LV transforms the coordinate system to account for growth rates, in addition to interaction terms (compare blue, orange, green *versus* purple coordinate systems). The red star in (A-D) marks an interaction matrix that is a generalized Prisoner’s Dilemma game when both growth rates are equal (A); see Fig. 2 for a full analysis of this example. (Parameters: *a*_11_ = −3, *a*_22_ = −4, *r*_1_ = *r*_2_ = 1 in A, *r*_1_ = 1.5 and *r*_2_ = 1 in B, *r*_1_ = 1 and *r*_2_ = −1 in C, and *r*_1_ = −0.5 and *r*_2_ = −1 in C; Red star: *a*_12_ = −5 and *a*_21_ = 1).

When *r*_1_ = *r*_2_, as we discussed above, the replicator dynamics Eq. (4) is dynamically equivalent to the Lotka-Volterra system and thus, dynamical outcomes of LV are accurately captured by the game-type classification [26–28] (shown by the purple plot in Fig. 1A): dominance of either type 1 or type 2, coexistence, or bistability. Because we assumed that *a*_11_ *> a*_22_, when the types are strategies and type 2 dominates it can be seen as a social dilemma (e.g., Prisoner’s Dilemma); when type 1 dominates, it is sometimes called ‘harmony’. Ecologically, this agrees with the fact that all mutualisms, some predator-prey interactions, and some competitive interactions are expected to lead to coexistence of the two species [25]. The remaining predator-prey interactions can lead only to dominance, i.e. the predator drives the prey extinct; and the remaining competitive interactions lead to bistability. Conversely, this overlaying of ecological and game theoretic worldviews also reveals that a Prisoner’s Dilemma is ecologically a predator-prey (parasitic) or competitive interaction, a coordination game is a competitive interaction, while a coexistence game (such as Hawk-Dove or Snowdrift) can be any type of ecological interaction—mutualistic, competitive, or predator-prey—depending on the sign of the payoffs that each type gets from the other. Importantly, though mutualisms have occasionally been discussed as the species-level analog of within-species cooperative interactions in the sense of game theory [29–33], this analysis shows that their appropriate analog are, in fact, coexistence games.

When *r*_1_ ≠ *r*_2_, writing the replicator dynamics solely based on the interaction matrix, and without regard to the difference in intrinsic growth rates, fails to accurately predict the dynamics of the two-type Lotka-Volterra system. This can be seen by comparing the purple system of coordinates against the blue one in Fig 1B, the orange one in Fig. 1C, and the green one in Fig 1D: the purple system captures the dynamics of (4), as in Fig. 1A, whereas the other three coordinate systems capture the dynamics of the Lotka-Volterra system with the same interaction matrix but accounting for *r*_1_ ≠ *r*_2_. When *r*_1_ *> r*_2_ *>* 0 (blue in Fig 1B), the region of coexistence is shifted up and to the left, so that, for example, more cases of predator-prey interactions that used to result in the extinction of species 1 (prey) now yield coexistence (compare the red star in Figs. 1A and B). If *r*_1_ *>* 0 *> r*_2_ (orange classification in Fig 1C) or 0 *> r*_1_ *> r*_2_ (green in Fig 1D), the game theoretic analysis gets wrong not just the precise regions corresponding to each possible outcome, but also the set of possible outcomes. Specifically, when *r*_1_ *>* 0 *> r*_2_ only coexistence or dominance of type 1 are possible (but not bistability or dominance of type 2), while when 0 *> r*_1_ *> r*_2_ only extinction is possible.

Thus, when *r*_1_ ≠ *r*_2_, there’s bad news and good news. The bad news is that one cannot apply the game-type classification and associated predictions and terminology from evolutionary game theory. The good news is that the shift in—and reclassification of—dynamical outcomes induced by *r*_1_ ≠ *r*_2_ is simple and elegant in its own right. For instance, if *r*_1_ *> r*_2_ *>* 0, then the horizontal line gets moved vertically from *a*_11_ to *a*_11_(*r*_2_*/r*_1_) and the vertical line gets moved horizontally from *a*_22_ to *a*_22_(*r*_1_*/r*_2_). Thus, coexistence occurs if *a*_21_ *> a*_11_(*r*_2_*/r*_1_) and *a*_12_ *> a*_22_(*r*_1_*/r*_2_), bistability occurs when both inequalities are reversed, and dominance occurs when the two inequalities have different signs (Fig. 1B blue). When *r*_1_ = *r*_2_, we recover the purple partitioning produced by the replicator dynamics (Fig. 1A). This graphical representation further allows us to see that changing the intrinsic growth rates relative to each other allows one to transform the dynamical outcome of any game. For instance, dominance of type 2 can be transformed into coexistence, bistability, or even into dominance of type 1, by increasing the growth rate of type 1 relative to type 2 (see SI Fig. C.1 for a summary of all possible transformations).

### Ceci n’est pas un… Prisoner’s Dilemma

To exemplify our results with a case study that has been amply studied both empirically and theoretically, we next focus on a generalized Prisoner’s Dilemma scenario, which is characterized by the payoff ranking *a*_21_ *> a*_11_ *> a*_22_ *> a*_12_. The dilemma is that type 2 is always better off in any pairwise interaction (owing to *a*_21_ *> a*_11_ and *a*_22_ *> a*_12_), but a homogeneous population of type 1 is better off than a homogeneous population of type 2 (owing to *a*_11_ *> a*_22_). Thus type 1 are the cooperators and type 2 are the defectors or free-riders.

The replicator equation (4) for the dynamics of the frequency of cooperators reveals that cooperators will always decrease in abundance, and that *x*_1_ = 0 (all defectors) is the only stable fixed point (see Fig 2A top). If *r*_1_ = *r*_2_, the Lotka-Volterra equations predict the same outcome, with the sole stable equilibrium being the defectors-only one (see Fig 2A bottom). If, however, the two types have different but positive growth rates, we can use the graphical illustration in Fig. 1B to predict the possible outcomes. Specifically, if *r*_1_ *> r*_2_ *>* 0 the outcome of a Prisoner’s Dilemma matrix can change from dominance to coexistence, provided that *a*_12_ *> a*_22_(*r*_1_*/r*_2_). Without loss of generality, we can assume *r*_2_ = 1. Keeping in mind that *a*_22_ *<* 0, the condition *a*_12_ *> a*_22_(*r*_1_*/r*_2_) is equivalent to *r*_1_ *> a*_12_*/a*_22_. For example, if the interaction matrix is *a*_11_ = − 3, *a*_12_ = − 5, *a*_21_ = +1 and *a*_22_ = − 4 (corresponding to the red star in Fig. 1), the above analysis predicts that the two types will coexist provided that *r*_1_ *>* 5*/*4. In other words, cooperators and defectors can coexist, as long as the intrinsic growth rate of cooperators is 25% greater than that of the defectors. This is, indeed, exactly what we see in Fig. 2B. In this case, although the matrix obeys the inequalities of a Prisoner’s Dilemma, the system is not trapped in a dilemma at all. We argue that, in this case, it would be misleading to continue to call the two types cooperators and defectors, as this nomenclature is immediately evocative of the game theoretic framework that ceases to apply when the two growth rates are not identical.

**Figure 2:**
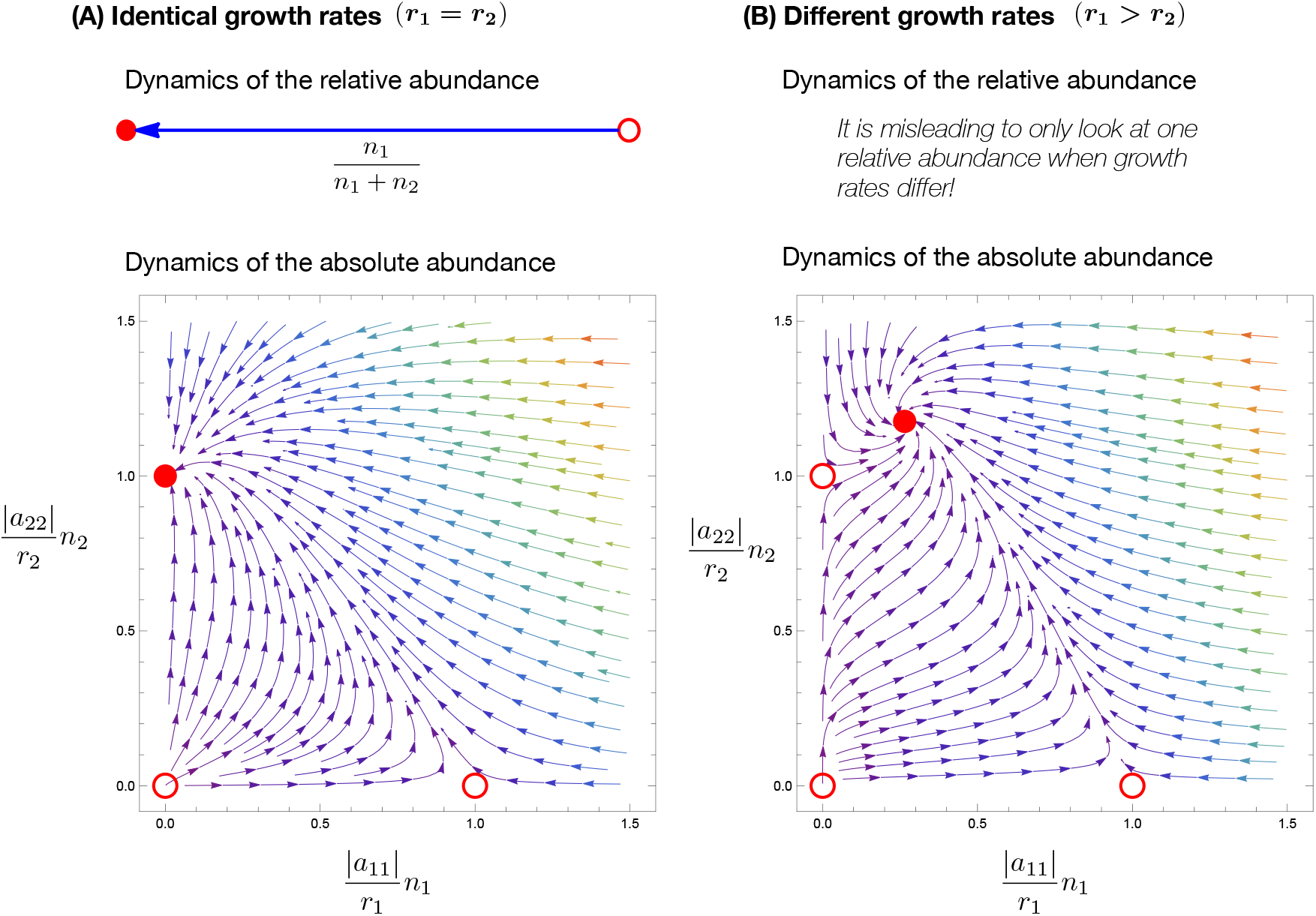
The dynamics of relative and absolute abundances for a general Prisoner’s Dilemma matrix. **(A)** When growth rates are identical, *r*_1_ = *r*_2_, the relative abundance is sufficient to capture the dynamics. Here, strategy 1 is cooperation and strategy 2 is defection. The only stable equilibrium (red disk) is defection: *x*_1_ = 0, *x*_2_ = 1 for relative abundances, and the corresponding (*n*_1_, *n*_2_) = (0, − *r*_2_*/a*_22_) for absolute abundances. Unstable equilibria are marked by red circles. **(B)** When growth rates are different, *r*_1_ = 2 and *r*_2_ = 1, looking at relative abundances alone is insufficient. In this case, the dynamics has three unstable fixed points at the boundaries, and a new and stable fixed point emerges in the interior. The term Prisoner’s Dilemma has no meaning anymore. Interaction parameters correspond to the red star in Fig. 1: *a*_11_ = −3, *a*_12_ = −5, *a*_21_ = +1, and *a*_22_ = −4.

### Assembling diverse communities, step-by-step

This is, in principle, good news for the study of behavior. Differential growth rates provide another simple mechanism for explaining why costly behaviors that are beneficial to others are apparently so widespread in nature. And the positive implications of this result extend beyond social behaviors, and turn out to be fundamental in the realm of ecology as well. Specifically, the realization that different growth rates can transform apparent dominance outcomes into coexistence of types unlocks an enormous potential for conceptualizing the assembly of ecologically complex communities. For example, cyclical dynamics have been occasionally found to exist empirically [5, 34, 35], despite skepticism from theoreticians who have pointed out that it is very hard for such cyclical interactions to be assembled, one mutant (or one migrant) at a time [36–38]. Intuitively, that is because cyclical interactions exist by definition only if all types are present; if one of the types is not present yet, then the broken cycle is a dominance hierarchy that leads to competitive exclusion. However, this intuition only holds when all growth rates are identical and the dynamical outcome is entirely predicted by the replicator equation. When the intrinsic growth rates of at least some of the types are different, the apparent dominance hierarchies can disappear and coexistence could be built step-by-step. In Appendix D and Figs. 3 and D.1 we exemplify how this would work in the simplest case of cyclical dynamics, that of Rock-Paper-Scissors.

**Figure 3:**
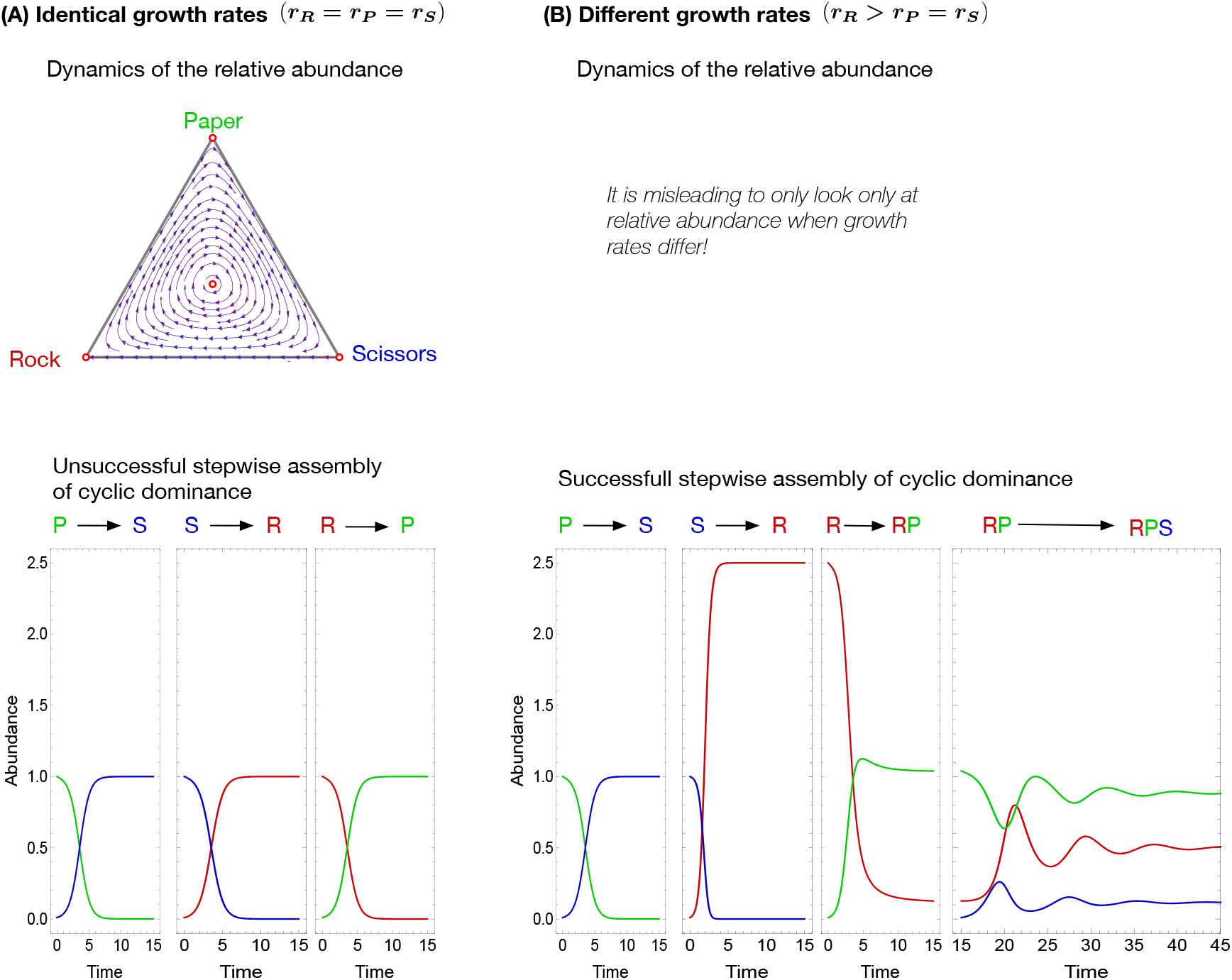
Assembling cyclic dynamics. We work with a fully symmetric Rock-Paper-Scissors (RPS) matrix, where winning leads to +*α* and losing to − *α* (see Appendix D). **(A)** (Top) For identical growth rates, the dynamics of the relative abundance shows cyclic motion on closed trajectories around a neutrally stable fixed point. (Bottom) One strategy can invade one other and be invaded by the third, such that a stepwise assembly of cyclic dominance via coexistence is not possible (bottom) if mutants (or migrants) are rare. **(B)** (Top) For asymmetric growth rates, it is misleading to consider the dynamics of relative abundances. (Bottom) A stepwise assembly via the coexistence of two strategies (in this case, *R* and *P*) into cyclic dominance of three types is possible. Interaction parameters: *a*_*ij*_ = − 1 *± α* with *α* = 1.3; growth rates: *r*_*R*_ = *r*_*P*_ = *r*_*S*_ = 1 in (A) and *r*_*P*_ = *r*_*S*_ = 1, *r*_*R*_ = 2.5 in (B).

This type of reasoning can be expanded to explore the assembly of cycles of any length, and even of more complex communities, such as those with higher order interactions. The assembly of such communities has remained a challenging open question despite active recent research—both theoretical [39, 40] and empirical [5, 41]—showing the potential of such interactions to maintain ecological diversity.

### So what does this all mean?

First and foremost, we argue that it is crucial for the field to emphatically acknowledge that the current setup of evolutionary game theory is limiting in terms of the biological systems that its predictions can apply to. Fundamentally, the framework of evolutionary game theory, from its inception in the 1970s, has been built upon the assumption that the payoff from the game is the sole source of differential fitness in the population. But this is, likely, rarely true in real systems. The field of evolutionary game theory has not reconsidered the implications of this original assumption, or made efforts to explicitly acknowledge it as a substantial caveat for its predictions, even as it moved to address other limitations, e.g., that the replicator dynamics ignores stochastic effects [21, 42, 43], that it assumes infinite population sizes [44], that it muddles growth rates and interactions [45, 46], or that it apparently ignores more realistic genetics and demography [47]. Where EGT applies, it remains a powerful framework, as originally demonstrated by the Hawk-Dove game and since then by many insights, especially for the study of social behavior in humans [48–50]. But, in non-human systems, and especially in microbes or cells, it should not be the default framework, at the very least not until intrinsic growth rates have been estimated. Otherwise there is high potential for paradoxical findings and unproductive debates.

Given the limitations of EGT, our paper adds further urgency to calls [51–53] for the field of social behavior to undergo a serious—albeit admittedly non-trivial—reconsideration of terminology. Terms such as cooperation, altruism, public good, cheater, or free-rider have become both unproductively anthropomorphized and inextricably linked with game theoretic predictions. But the framework itself very likely does not apply as broadly as one encounters systems with costly behaviors that provide some benefit to others. Take, for instance, invertase-production in the yeast *Saccharomyces cerevisiae*. In a sucrose medium, yeast produces invertase to hydrolyze the sucrose into glucose, a preferred nutrient. Because invertase is costly to produce and 99% of its benefits diffuse away before they can be imported into the producing cell [12]—and are thus freely available to others—invertase has been labeled a costly public good and invertase production a cooperative behavior [7, 54]. The cooperator label naturally led empiricists to invoke the predictions of evolutionary game theory and hypothesize that the outcome of a well-mixed interaction between producers and non-producers should be a tragedy of the commons, i.e. invertase-producers should be outcompeted by non-producers; instead, producers and non-producers were found to stably coexist in lab experiments [12]. The reason is that when invertase-producing yeast is rare and in a sucrose-only medium, its ability to privatize 1% of the glucose it transforms gives it a substantial intrinsic growth advantage over the knockout non-producing strain under identical conditions [12]. In other words, producers and non-producers do not have identical intrinsic growth rates in a sucrose-only medium. In this light, the coexistence outcome [12] is only puzzling and in dissonance with the theory because the theory does not directly apply to this system without giving consideration to the growth rate differences. If, instead, the departure point had been the Lotka-Volterra framework, the starting theoretical hypothesis would have been that the mix of the two strains will result either in tragedy of the commons or in coexistence, depending on how much higher the intrinsic growth rate of producers was than that of non-producers (Fig. 1B). The experimental results would then have naturally existed within the set of possible theoretical outcomes. This ultimately interesting study of diverse microbial consumer-resource interactions illustrates the pitfalls of a deceptively intuitive terminology (‘cooperation’ and ‘costly public goods’) evocative of a deceptively broad framework. Invertase production is by no means an isolated example; other costly extracellular products have been painted with the broad brush stroke of cooperation, public goods, and the tragedy of the commons when the reality is, in fact, more nuanced (see, e.g., iron scavenging and siderophore production [15, 52, 53], where there is, similarly, some degree of privatization).

Microbial experiments are also useful to illustrate our final takeaway. Both replicator dynamics and Lotka-Volterra are phenomenological in their description of how the cost is paid, how the benefit is produced, and how it is being used. When employed appropriately, this makes these frameworks good candidates for ‘first approximation’-predictions of what one might expect the outcome to be (e.g., either coexistence or dominance in the invertase-producing yeast scenario). However, when one fully controls the experimental setup (what goes in, what comes out etc.) and has a more mechanistic understanding of the process, then one could aim for a still simple but more system-specific theoretical model that allows for a more meaningful and in depth analysis [55]. For one, such an approach would address an existing criticism that most ‘cooperative’ microbial interactions are between many partners rather than between pairs and that, accordingly, employing predictions from pairwise games is fundamentally inadequate [56–58]. Of course, in empirical systems that are less tractable, ‘first approximation’-predictions will continue to play an important role and there it is imperative to employ the broader, ecological framework.

Although our focus here has been on the replicator dynamics in well-mixed populations, which is the bedrock of EGT, the same arguments apply to extensions of the framework to finite and/or structured populations, where the same original assumption of identical intrinsic growth rates has been embraced. Reconsidering the entire body of work of evolutionary game theory to account for intrinsic growth rates will likely lead to a productive rethinking of predictions, including in the realm of structured populations [27, 50]. Conversely, because evolutionary game theory has developed sophisticated methods for thinking about a variety of complex interactions that only recently have been proposed—but remain understudied—in ecology, reestablishing the connection with ecological theory could substantially inform the latter in the study of complex species interactions (e.g., higher-order, spatially structured, multiscale, hierarchical etc.).

## Acknowledgements

We are indebted to Simon Levin, Jonathan Levine, Joshua Plotkin, and members of the Theoretical Ecology group at Princeton and the Department of Theoretical Biology in Plön for many helpful comments at all stages of this work.

## A General number of types

For *m* types the Lotka-Volterra equations are

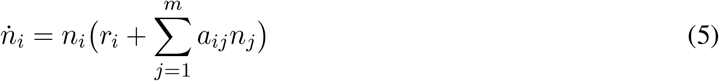

where *i* = 1, …, *m*. Then the frequency of individuals of type *i* is *x*_*i*_ = *n*_*i*_*/n*, where 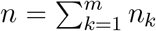 is the total population size. The dynamics of frequencies can be derived from the dynamics of abundances using the quotient rule:

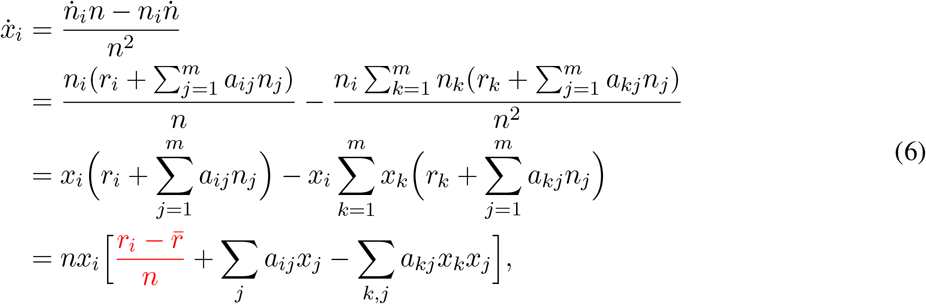

where 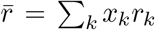 is the average intrinsic growth rate of the population and the two summation terms can be recognized as *f*_*i*_ = ∑_*j*_ *a*_*ij*_*x*_*j*_ (the fitness of type *i*) and 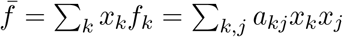 (the average fitness). Therefore, Eq. (6) can be rewritten as:

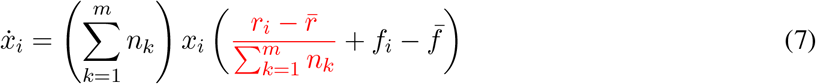

When *m* = 2, Eq. (7) yields Eq. (3). The term marked in red can be ignored only when the system is unbounded or the intrinsic growth rate of every type is equal to the average growth rate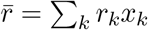, which is equivalent to all types having identical intrinsic growth rates.

## B Frequency-dependent ecological interactions

For *m* types, the abundance equations with frequency-dependent interaction effects are

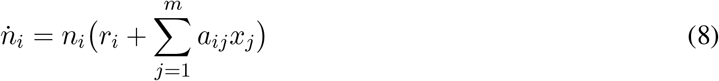

where *i* = 1, …, *m*. The frequency of individuals of type *i* can be derived as above using the quotient rule:

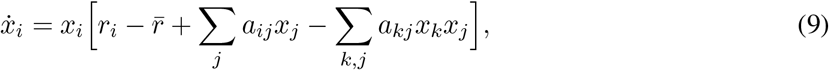

where 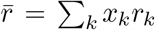 is the average intrinsic growth rate of the population and the two summation terms can be recognized as *f*_*i*_ = ∑_*j*_ *a*_*ij*_*x*_*j*_ (the fitness of type *i*) and 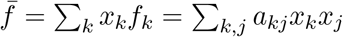 (the average fitness). Rewriting *r*_*i*_ = *r*_*i*_ ∑_*j*_ *x*_*j*_ as well as 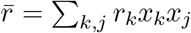 and combining each *a*_*ij*_ term with an *r*_*i*_, Eq. (9) becomes a replicator dynamics for the *m*-type game with payoff matrix Ã = [*r*_*i*_ + *a*_*ij*_]:

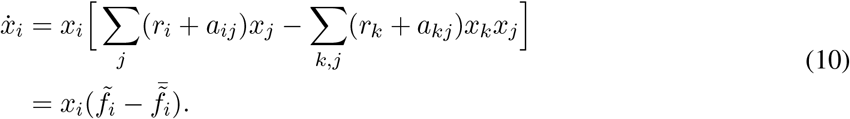

In other words, when the ecological interactions are frequency dependent, there is no ‘problematic’ term and there exists a direct transformation of the ecological system with *m* types into an *m*-type game whose matrix is modified to account for the payoffs. Although, at first glance, this might appear like good news, the fundamental problem of different intrinsic growth rates changing the outcome of the game remains. Specifically, let us for simplicity take the case *m* = 2: if one attempts an evolutionary game theoretical treatment for the *m*-type system with matrix *A*

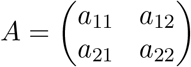

and thus, consistent with the setup of EGT, never asks oneself about the possibility of different intrinsic growth rates, then one would attempt to simply make predictions based on the matrix *A*. However, the correct predictions should be made for the transformed matrix *A*

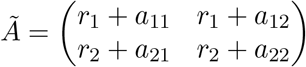

and these predictions will differ. For instance, the correct condition that strategy *A*_1_ is evolutionarily stable (ESS) would be *r*_1_ + *a*_11_ *> r*_2_ + *a*_21_, which is not equivalent to *a*_11_ *> a*_21_ when *r*_1_ ≠ *r*_2_.

## C Possible transitions

Using asymmetric growth rates, we can transform the game between different scenarios, cf. Fig. 1. Depending on the sign and the ranking of the payoffs, all possible transitions are shown in Fig. C.1.

**Figure C.1:**
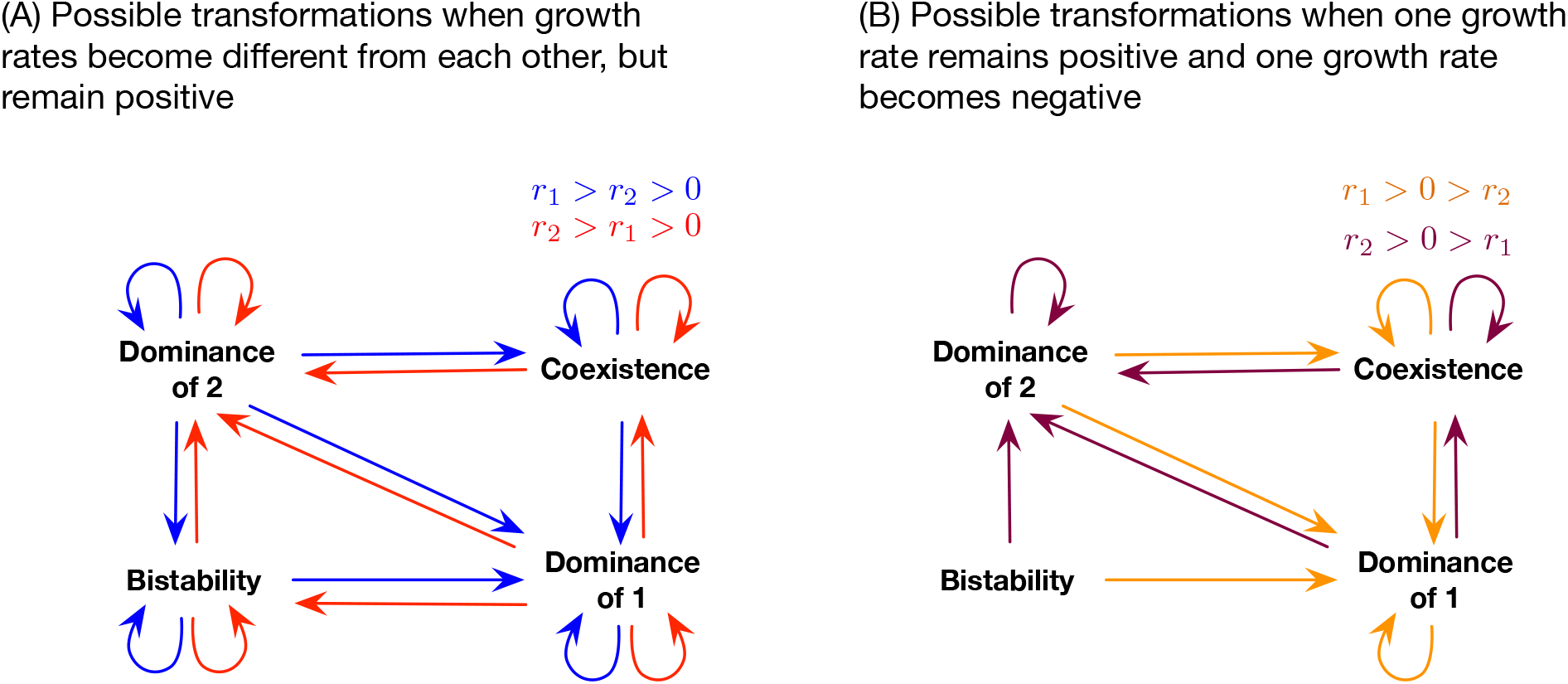
Possible transitions between stability scenarios. (A) Starting from symmetric growth rates, almost any stability transition can be achieved if one of the growth rates can be changed – even if growth rates remain positive. However, coexistences cannot become bistabilities (and vice versa). (B) When one growth rate becomes negative, the bi-stabilities vanish entirely and some dominances vanish as well.

## D Step-wise assembly of a Rock-Paper-Scissors dynamics

We consider the simplest RPS matrix (i.e. completely symmetric, having a single parameter *α >* 0) that also incorporates the assumption of bounded growth for each individual type:

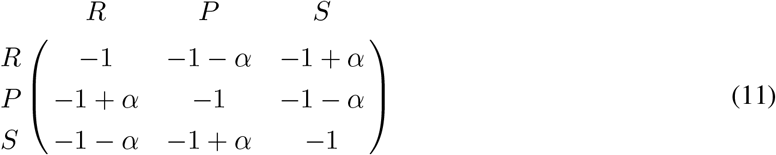

Writing the 3-type Lotka-Volterra equations for the dynamics of their abundances, we obtain

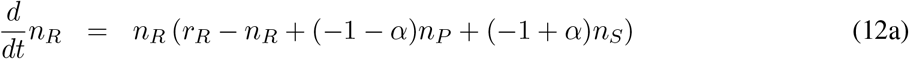

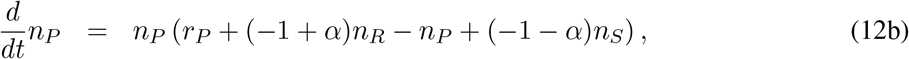

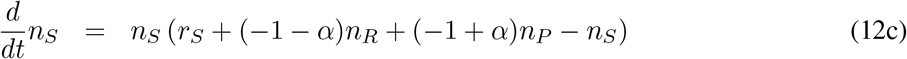

where *r*_*R*_, *r*_*P*_, and *r*_*S*_ are the intrinsic growth rates of *R, P*, and *S*. The fixed points of this dynamics are

- Extinction of the population, (*n*_*R*_, *n*_*P*_, *n*_*S*_) = (0, 0, 0). Assuming positive growth rates, this fixed point is unstable
- Homogeneous populations, (*n*_*R*_, *n*_*P*_, *n*_*S*_) = (*r*_*R*_, 0, 0), (*n*_*R*_, *n*_*P*_, *n*_*S*_) = (0, *r*_*P*_, 0), (*n*_*R*_, *n*_*P*_, *n*_*S*_) = (0, 0, *r*_*S*_). Positive growth rates ensure that these three fixed points are saddle points.
- Coexistence of two types (if the fixed point coordinates are positive).

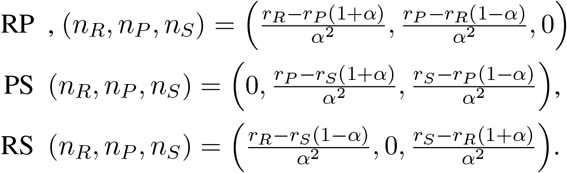
- Finally, there is a fixed point RPS where all three types coexist (if its coordinates are positive)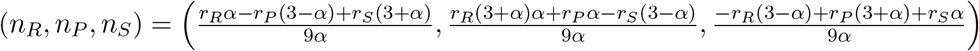.

We first confirm that for *r* = *r*_*R*_ = *r*_*P*_ = *r*_*S*_ *>* 0 the dynamics has unstable fixed points at (*r*, 0, 0), (0, *r*, 0), and (0, 0, *r*). In addition, there is—for symmetry reasons—a neutrally stable fixed point at (*r/*3, *r/*3, *r/*3). Thus, when the three growth rates are the same, no two types can coexist but all three will coexist and cycle around the neutrally stable fixed point [59]. The question is: Can this internal equilibrium be assembled pairwise, one mutant at a time, if not all growth rates are equal?

To answer this, we pick one of the three pairwise interactions, e.g., between Rock and Paper, and transform the outcome of the game, which would otherwise result in dominance of Paper, into coexistence by changing the growth rate *r*_*R*_ similar to the graphical illustration in Fig. 1B. We fix *r*_*P*_ = *r*_*S*_ = *r >* 0 and let *r*_*R*_ = *r* + *ρ*. To destabilize Paper, we need *r*_*R*_ *>* (1 + *α*)*r*_*P*_, i.e. *ρ/r > α*. In addition, we need to ensure that Rock alone remains unstable so that we can get coexistence, which is the case if either *α* ≥ 1 or if *α <* 1 and *ρ/r < α/*(1 − *α*). With growth rates fulfilling these conditions, now Rock and Paper can coexist stably in the absence of Scissors at equilibrium abundances 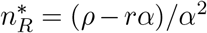 and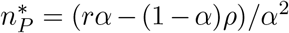. Once Scissors arrives, either via mutation or via immigration, its initial growth rate is 3*r* + *ρ* − 3*ρ/α*. Scissors can invade only if this growth rate is positive, i.e. either if *α* ≥ 3 or if *α <* 3 and *ρ/r <* 3*α/*(3 − *α*). Note that this last condition is always stronger than the condition to keep Rock unstable. Thus, the parameter range in which Scissors can invade the coexistence of Rock and Paper is narrower than the parameter range for a stable coexistence of Rock and Paper. The scenario of a stable coexistence and the possible invasion of the third type—holds for 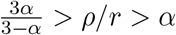 or *ρ/r > α >* 3. The table below summarizes these conditions.

| Condition | $0 < \alpha < 1$ | $1 < \alpha < 3$ | $3 < \alpha$ |
| --- | --- | --- | --- |
| RP is stable | $\alpha < \frac{\rho}{r} < \frac{\alpha}{1-\alpha}$ | $\alpha < \frac{\rho}{r}$ | $\alpha < \frac{\rho}{r}$ |
| RPS is stable | $\frac{\rho}{r} < \frac{3\alpha}{3-\alpha}$ | $\frac{\rho}{r} < \frac{3\alpha}{3-\alpha}$ | for all $\frac{\rho}{r}$ |

The only remaining questions are whether this newly assembled system still leads to stable coexistence of the three types and whether it still does so in a manner that reflects cyclical interactions. Specifically, the coexistence point needs to be either neutrally stable or stable and approached dynamically via dampened oscillations. A numerical sweep of the eigenvalues of the coexistence point confirmed the above conditions for the assembly, as well as the stability of the fixed point (Fig. D.1). Moreover, we have also confirmed numerically that the imaginary eigenvalues have a negative real part when the equilibrium is stable and, thus, that the stable equilibrium is approached via dampened oscillations (Fig. D.1).

**Figure D.1:**
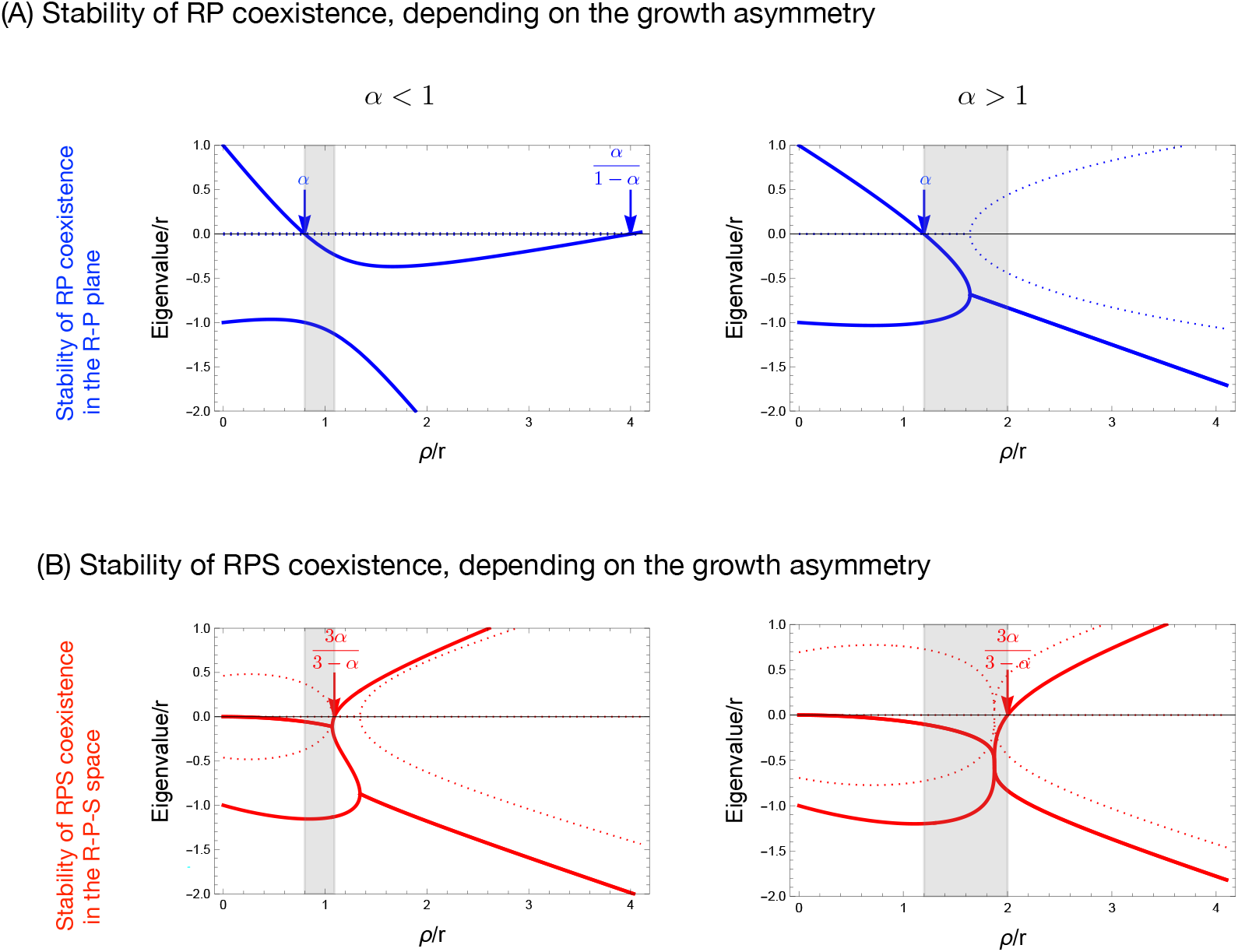
Stability in the rock paper scissors dynamics with asymmetric growth rates. We show a linear stability analysis of the fixed points with Rock and Paper only (A) and with Rock, Paper and Scissors (B), focusing on the Eigenvalues of the linearized system at the fixed points. Real parts of Eigenvalues are full lines, imaginary parts dotted lines. If all Eigenvalues have negative real parts, the corresponding fixed point is stable. Regardless of whether *α <* 1 or *α >* 1, there exists a subregion (marked in gray) for 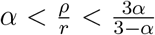 where Rock and Paper stably coexist, Scissors can invade, and the dynamics spirals into the RPS fixed point (Parameters *r* = 1, *ρ* = 1.5, *α* = 0.8 in (A) and *α* = 1.3 in (B)).

## Footnotes

* A slightly different ecological approach consists of writing similar abundance equations but where the interaction effects are frequency-dependent, i.e.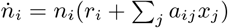. We will address it briefly below and in Appendix B.

† Note that, although *a priori* it might seem like the interaction terms in a Lotka-Volterra system should have different units than those in the replicator dynamics (specifically, 1*/*(*t ×* number) versus 1*/t*), the non-dimensionalization is accounted for by the dynamical rescaling term *n*_1_ + *n*_2_ *>* 0, so that equations (2) and (4) are, indeed, written for the same matrix *A*.

‡ In fact, Hofbauer and Sigmund [22] demonstrated that the replicator dynamics for an **m-type game** is equivalent to the generalized Lotka-Volterra dynamics for an **(m-1)-type ecological system** when the intrinsic growth rates are different (e.g., a Prisoner’s Dilemma is equivalent to a 1-species population growth but not to a general predator-prey interaction).

§ This remains true even if the ecological interactions are frequency, rather than density, dependent (see Appendix B).

## References

1. Maynard Smith, J. & Price, G. R. The logic of animal conflict. Nature 246, 15–18 (1973).

2. Taylor, P. D. & Jonker, L. B. Evolutionarily stable strategies and game dynamics. Mathematical Biosciences 40, 145–156 (1978).

3. Zeeman, E. C. Population dynamics from game theory. Lecture Notes in Mathematics 819, 471–497 (1980).

4. Velicer, G. J., Kroos, L. & Lenski, R. E. Loss of social behaviors by Myxococcus xanthus during evolution in an unstructured habitat. Proceedings of the National Academy of Sciences USA 95, 12376–12380 (Feb. 1998).

5. Kerr, B., Riley, M. A., Feldman, M. W. & Bohannan, B. J. M. Local dispersal promotes biodiversity in a real-life game of rock-paper-scissors. Nature 418, 171–174 (2002).

6. Rainey, P. B. & Rainey, K. Evolution of cooperation and conflict in experimental bacterial populations. Nature 425, 72–74 (2003).

7. Greig, D. & Travisano, M. The Prisoner’s Dilemma and polymorphism in yeast SUC genes. Proceedings of the Royal Society of London B: Biological Sciences 271, S25–S26 (2004).

8. Griffin, A. S., West, S. A. & Buckling, A. Cooperation and competition in pathogenic bacteria. Nature 430, 1024–1027 (2004).

9. Tomlinson, I. Game-Theory Models of Interactions between Tumour Cells. European Journal of Cancer 33, 1495–1500 (Aug. 1997).

10. Archetti, M., Ferraro, D. A. & Christofori, G. Heterogeneity for IGF-II production maintained by public goods dynamics in neuroendocrine pancreatic cancer. Proceedings of the National Academy of Sciences 112, 1833–1838 (2015).

11. Kaznatcheev, A., Peacock, J., Basanta, D., Marusyk, A. & Scott, J. G. Fibroblasts and Alectinib Switch the Evolutionary Games Played by Non-Small Cell Lung Cancer. Nature Ecology & Evolution 3, 450–456 (Feb. 2019).

12. Gore, J., Youk, H. & van Oudenaarden, A. Snowdrift game dynamics and facultative cheating in yeast. Nature 459, 253–256 (2009).

13. Van Dyken, J. D., Müller, M. J. I., Mack, K. M. L. & Desai, M. M. Spatial population expansion promotes the evolution of cooperation in an experimental Prisoner’s Dilemma. Current biology 23, 919–923 (2013).

14. Zhang, X.-X. & Rainey, P. B. Exploring the sociobiology of pyoverdin-producing Pseudomonas. Evolution 67, 3161–3174 (2013).

15. Kümmerli, R. & Ross-Gillespie, A. Explaining the sociobiology of pyoverdin producing Pseudomonas: a comment on Zhang and Rainey (2013). Evolution 68, 3337–3343 (2014).

16. Von Neumann, J. & Morgenstern, O. Theory of Games and Economic Behavior 3rd (Princeton University Press, Princeton, 1953).

17. Lewontin, R. C. Evolution and the theory of games. Journal of Theoretical Biology 1, 382–403 (1961).

18. Slobodkin, L. B. The strategy of evolution. American Scientist 52, 342–357 (1964).

19. Schuster, P. & Sigmund, K. Replicator dynamics. Journal of Theoretical Biology 100, 533–538 (1983).

20. Fisher, R. A. in The Genetical Theory of Natural Selection chap. Sexual Reproduction and Sexual Selection (Clarendon Press, Oxford, 1930).

21. Wang, G., Su, Q., Wang, L. & Plotkin, J. B. Reproductive variance can drive behavioral dynamics. Proceedings of the National Academy of Sciences 120, e2216218120 (2023).

22. Hofbauer, J. & Sigmund, K. Evolutionstheorie und dynamische Systeme – Mathematische Aspekte der Selektion (Verlag Paul Parey, Berlin, Hamburg, 1984).

23. Bomze, I. M. Lotka-Volterra equation and replicator dynamics: a two-dimensional classification. Biological Cybernetics 48, 201–211 (1983).

24. Cressman, R. in (Springer Verlag, 1992).

25. Mittelbach, G. G. & McGill, B. J. Community Ecology (Oxford University Press, 2019).

26. Hauert, C. Effects of space in 2 2 Games. International Journal of Bifurcation and Chaos 12, 1531– 1548 (2002).

27. Szabó, G. & Fáth, G. Evolutionary games on graphs. Physics Reports 446, 97–216 (2007).

28. Nowak, M. A. Evolutionary dynamics: Exploring the equations of life (Harvard University Press, 2006).

29. Shou, W., Ram, S. & Vilar, J. M. G. Synthetic cooperation in engineered yeast populations. Proceedings of the National Academy of Sciences of the United States of America 104, 1877–1882 (2007).

30. Momeni, B., Brileya, K. A., Fields, M. W. & Shou, W. Strong inter-population cooperation leads to partner intermixing in microbial communities. Elife 2 (2013).

31. Foster, K. R. & Bell, T. Competition, not cooperation, dominates interactions among culturable microbial species. Current Biology 22, 1845–1850 (2012).

32. Coyte, K. Z., Schluter, J. & Foster, K. R. The ecology of the microbiome: Networks, competition, and stability. Science 350, 663–666 (2015).

33. Pauli, B., Oña, L., Hermann, M. & Kost, C. Obligate mutualistic cooperation limits evolvability. Nature Communications 13, 337 (2022).

34. Sinervo, B. & Lively, C. M. The rock-paper-scissors game and the evolution of alternative male strategies. Nature 380, 240–243 (1996).

35. Kirkup, B. C. & Riley, M. A. Antibiotic-mediated antagonism leads to a bacterial game of rock–paper– scissors in vivo. Nature 428, 412–414 (Mar. 2004).

36. Reichenbach, T., Mobilia, M. & Frey, E. Mobility promotes and jeopardizes biodiversity in rock–paper– scissors games. Nature 448, 1046–1049 (2007).

37. Park, H.-J., Pichugin, Y. & Traulsen, A. Why is cyclic dominance so rare? eLife 9, 10.7554/eLife.57857 (2020).

38. Vandermeer, J. & Perfecto, I. Intransitivity as a dynamic assembly engine of competitive communities. Proceedings of the National Academy of Sciences 120, e2217372120 (2023).

39. Levine, J. M., Bascompte, J., Adler, P. B. & Allesina, S. Beyond pairwise mechanisms of species coexistence in complex communities. Nature 546, 56–64 (2017).

40. Kotil, S. E. & Vetsigian, K. Emergence of evolutionarily stable communities through eco-evolutionary tunnelling. Nature Ecology & Evolution 2, 1644–1653 (2018).

41. Kelsic, E. D., Zhao, J., Vetsigian, K. & Kishony, R. Counteraction of antibiotic production and degradation stabilizes microbial communities. Nature 521, 516–519 (2015).

42. Traulsen, A., Claussen, J. C. & Hauert, C. Coevolutionary dynamics: From finite to infinite populations. Physical Review Letters 95, 238701 (2005).

43. Constable, G. W. A., Rogers, T., McKane, A. J. & Tarnita, C. E. Demographic noise can reverse the direction of deterministic selection. Proceedings of the National Academy of Sciences 113, E4745– E4754 (2016).

44. Nowak, M. A., Sasaki, A., Taylor, C. & Fudenberg, D. Emergence of cooperation and evolutionary stability in finite populations. Nature 428, 646–650 (2004).

45. Gerlee, P. & Altrock, P. M. Complexity and stability in growing cancer cell populations. Proceeding of the National Academy of Sciences 112, 2742–2743 (2015).

46. Yamamichi, M., Tsuji, K., Sakai, S. & Svensson, E. I. Frequency-dependent community dynamics driven by sexual interactions. Population Ecology 65, 204–219 (2023).

47. Van Cleve, J. Evolutionarily stable strategy analysis and its links to demography and genetics through invasion fitness. Philosophical transactions of the Royal Society B 378, 20210496 (2023).

48. Nowak, M. A. Five rules for the Evolution of Cooperation. Science 314, 1560–1563 (2006).

49. Yoeli, E. & Hoffman, M. Hidden Games: The Surprising Power of Game Theory to Explain Irrational Human Behavior (Basic Books, 2022).

50. Traulsen, A. & Glynatsi, N. The future of theoretical evolutionary game theory. Philosophical Transactions of the Royal Society B 378, 20210508 (2023).

51. Tarnita, C. E. The ecology and evolution of social behavior in microbes. Journal of Experimental Biology 220, 18–24 (2017).

52. Zhang, X.-X. & Rainey, P. B. Exploring the sociobiology of pyoverdin-producing Pseudomonas. Evolution 67, 3161–3174 (2013).

53. Rainey, P. B., Desprat, N., Driscoll, W. W. & Zhang, X.-X. Microbes are not bound by sociobiology: Response to Kümmerli and Ross-Gillespie (2013). Evolution (2014).

54. MacLean, G., Fuentes-Hernandez, A., Greig, D., Hurst, L. D. & Gudelj, I. A mixture of “cheats” and “co-operators” can enable maximal group benefit. PLoS Biology 8, e1000486 (2010).

55. Pfeiffer, T., Schuster, S. & Bonhoeffer, S. Cooperation and Competition in the Evolution of ATP-Producing Pathways. Science 292, 504–507 (Aug. 2001).

56. Hauert, C., Michor, F., Nowak, M. A. & Doebeli, M. Synergy and discounting of cooperation in social dilemmas. Journal of Theoretical Biology 239, 195–202 (2006).

57. Pacheco, J. M., Santos, F. C., Souza, M. O. & Skyrms, B. Evolutionary dynamics of collective action in N-person stag hunt dilemmas. Proceedings of the Royal Society B 276, 315–321 (2009).

58. Peña, J., Lehmann, L. & Nöldeke, G. Gains from switching and evolutionary stability in multi-player matrix games. Journal of Theoretical Biology 346, 23–33 (2014).

59. May, R. M. & Leonard, W. J. Nonlinear Aspects of Competition Between Three Species. SIAM Journal on Applied Mathematics 29, 243–253 (June 1975).

